# Weighted Ensemble Approach for Knowledge Graph completion improves performance

**DOI:** 10.1101/2024.07.16.603664

**Authors:** Meghamala Sinha, Roger Tu, Carolina González, Andrew I. Su

## Abstract

This study introduces a weighted ensemble method called “WeightedKgBlend” for link prediction in knowledge graphs which combines the predictive capabilities of two types of Knowledge Graph completion methods: knowledge graph embedding and path based reasoning. By dynamically assigning weights based on individual model performance, WeightedKgBlend surpasses standalone methods, demonstrating superior predictive accuracy when tested to discover drug-disease candidates over a large-scale biomedical knowledge graph called Mechanistic Repositioning Network. This research highlights the efficacy of an integrated approach combining multiple methods in drug discovery, showcasing improved performance and the potential for transformative insights in the realm of biomedical knowledge discovery.

## Introduction

In the ever-evolving field of drug discovery, knowledge graph completion has proven to be extremely useful due to its potential to uncover new insights and accelerate the development of new treatments. By predicting missing links and inferring new relationships within complex biomedical data, knowledge graph completion methods helps identify novel drug-target interactions, repurpose existing drugs, predict adverse reactions, understand disease mechanisms, and discover new biomarkers, ultimately contributing to more effective and personalized treatments. Numerous knowledge graph completion methods^1^ have employed efficient and novel mechanism to predict missing connections or to deduce new relationships, each with their own strength. However seldom work has been done to combine their potency together. Herein lies the essence of ensemble methods, a powerful strategy that harnesses the collective intelligence of diverse computational models to enhance individual method’s predictive accuracy and robustness. In this research we aim to create a comprehensive and synergistic approach to drug discovery by integrating two solid methods: knowledge graph embedding (KGE)^2^ and a type of path based reasoning method known as case-based reasoning (CBR)^3^. Furthermore, we also show our method’s potential applicability for Drug Repurposing^4^, where knowledge graphs can integrate information about drug properties, mechanisms of action, and clinical outcomes from existing data and the algorithm can identify existing drugs that could be repurposed for new therapeutic uses based on their similarity to other drugs or diseases in the graph.

KGE based methods are perfect for capturing complex relationships in structured data like biomedical knowledge graphs. These methods provide powerful tools for modeling complex biological interactions by representing entities and relationships as vectors in continuous vector space. On the other hand, CBR methods leverage historical cases to make predictions or decisions in new, analogous situations. It represents knowledge in the form of a case, each containing problems, solutions and successful solution contexts. CBR systems are generally more interpretable, as solutions are derived from specific cases. The reasoning process can be explained by referencing past instances. So CBR method like path based reasoning methods help analyzing paths or sequences of relations between entities in the knowledge graph.

Recognizing the complementary strengths of KGE and CBR, our proposed approach introduces a weighted ensemble method called “WeightedKgBlend” that fuses the predictive capabilities of various embedding algorithms and CBR model. By assigning weights dynamically based on the performance of each individual method, our ensemble method aims to capitalize on the unique strengths of each technique while mitigating their respective limitations. WeightedKgBlend aims to improve the accuracy and robustness of drug prediction and offers promising opportunities to discover new therapeutic uses of existing drugs. This method has been tested on Mechanistic Repositioning Network (MRN), a large heterogeneous network with an widely expressed relationships describing a drug’s mechanism of action^5^.

The main mechanism of WeightedKgBlend is a simple ensemble technique that combines predictions from multiple models, in our case five KGE methods and two CBR methods, assigning different weights to each model’s prediction based on its performance. This typically works by first training the individual models followed by generating predictions by each model. Since, we are predicting [drug→ treats→ disease], the predictions are a list of ranked diseases (other than the expected indicated disease) that represents potential repurposing targets of the drug. Next, weights are assigned to these individual predictions and optimised based on performance, models with better performance may receive higher weights, while poorer-performing models may receive lower weights. Finally, the predictions of all the models are combined using the assigned weights to produce the final ensemble prediction. We evaluate performance of the final ensemble based on the typical train/ test/ valid split and report WeightedKgBlend to perform superior than all the individual methods across various performance metrics The following sections explore in details the theoretical basis, methodology, and experimental analysis of our proposed method and highlight the potential to leverage the strengths of multiple models while mitigating their weaknesses for the research and development of drug discovery.

## Related work

### 0.1 Knowledge Graph completions methods

Knowledge graph embedding (KGE) methods have significantly contributed to the field of knowledge graph completion by providing efficient and effective representations of entities and relations, enabling various downstream tasks such as link prediction and entity recommendation. Here is a brief description of the different types of KGE methods which we use for our method.

TransE: Translating Embeddings for Modeling Multi-relational Data (TransE) is a knowledge graph embedding model^6^. It represents entities and relations as low-dimensional vectors in the same space and models relations as translations operating on entity embeddings. Specifically, TransE aims to minimize the energy of correct triples while maximizing the margin with respect to incorrect triples.

RotatE: RotatE^7^ extends the idea of knowledge graph embeddings by representing relations as rotations in complex vector space. Unlike TransE, which assumes linear translations, RotatE captures the rotational patterns inherent in some relations. It achieves state-of-the-art performance on knowledge graph completion tasks by modeling relation-specific rotations.

DistMult: DistMult^8^ is a simple yet effective knowledge graph embedding model that extends the bilinear model for relation prediction. It represents entities and relations as low-dimensional vectors and computes scores for triples using the Hadamard product (element-wise multiplication) between entity embeddings. DistMult has shown competitive performance on link prediction tasks.

ComplEx: ComplEx^9^ extends the DistMult model by representing relations as complex-valued vectors. By incorporating complex-valued embeddings, ComplEx captures asymmetric relations and interactions between entities and relations more effectively than real-valued models like DistMult. ComplEx has demonstrated superior performance on knowledge graph completion tasks.

Apart from the embedding models, we also explored Rephetio ^10^, short for “repositioning drugs by integrating heterogeneous data,” integrates diverse data sources, including genomic, chemical, phenotypic, and clinical data, into a unified computational framework to predict potential drug-disease associations. Rephetio employs a machine learning approach that learns from known drug-disease associations to make predictions about novel drug indications. By considering multiple layers of biological information and utilizing network-based algorithms, Rephetio offers a systematic and data-driven approach to drug repurposing.

Case-based reasoning (CBR) is a problem-solving approach in artificial intelligence that involves solving new problems based on past experiences, or “cases”. Instead of relying on explicit rules or generalizations, CBR systems use a retrieval-and-adaptation process to find similar cases from a case library and adapt them to address the current problem. This method is inspired by how humans often solve problems by recalling similar past experiences and applying them to new situations. The four main steps involved in CBR are namely: Retrieve, Reuse, Revise and Retain. We are using a simple CBR approach over Knowledge Bases^11^ considering the task of finding a target entity given a source entity and a binary relation. We are also using Prob-CBR^12^, an extension of the simple CBR method which learns to weight rules with respect to clusters of entities and query relation and is more efficient to handle updates in an open world KG settings.

Ensemble learning as an approach to combine predictions from several models with the goal of achieving a better performance than any of the individual models has been previously explored^13^ using a limited number of embedding methods available then^14,15^. More as of late, Choi et al. displayed a comparative concept in which rather than employing a subset of the triples, the whole distribution of triple scores is normalized to be utilized by a Product of Experts (PoE) to ensemble results^**?**^. Recent studies^16^ also extends the concept of ensembling popular embedding methods to improve predictions in biomedical KGs designed for drug discovery. While these are studies that explore embedding methods, seldom work has been done to include different types of KG completion methods like path-based methods to utilize their interpretability capabilities.

## Methodology

### 0.2 Dataset preparation

In this study, we evaluate our experiments on Mechanistic Repositioning Network (MRN), a comprehensive biomedical knowledge graph created by integrating 18 different data sources, composed of 9,652,116 edges, 250,035 nodes, 9 types of nodes, and 22 relationships^5,17^. To add more reliability in MRN, we map edges in MRN which are listed as indications approved by t he Food and Drug Administration (FDA). For this, we resort to Drugcentral^18^, an open-access online drug resource which integrates structure, bioactivity, regulatory, pharmacologic actions and indications approved by FDA and other regulatory agencies.

We were able to map 5558 indications from DrugCentral to MRN which were added to MRN as FDA approved and more reliable [drug→ treats→ disease] tuple. We then split these indications into a train (80%), test (10%), validation (10%) to test all our benchmark KG completion algorithms. We named this new MRN dataset mapped with indications as MIND^19^.

### 0.3 Our Approach

In this section, we test that ensembling KGE methods with CBR methods by combining predictions for link prediction can be a powerful approach to leverage the strengths of both techniques. As displayed in Fig 1, we do this in the following steps:

**Figure 1.**
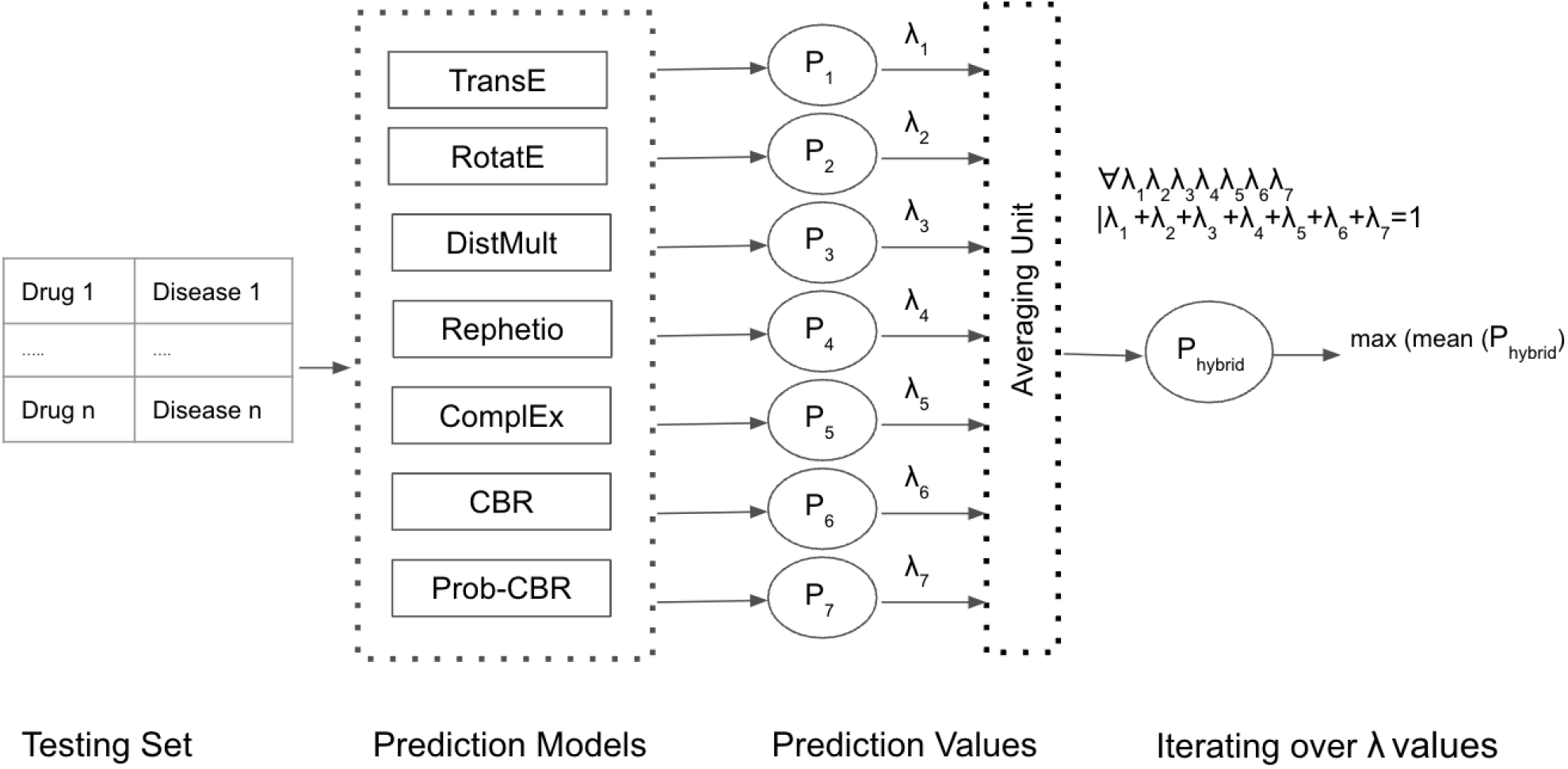
Visual representation of the workflow of the ensemble method: Given the prediction models, we find predictions for each test query consisting of [Drug→ treats→ Disease] triple. This results in a list of predicted diseases each with a corresponding score. Given these lists, we compute a weighted average for all possible combinations of (*λ*_1_, *λ*_2_, *λ*_3_, *λ*_4_, *λ*_5_, *λ*_6_, *λ*_7_) to find the combination resulting in the highest combination score.

- Data Preparation: First we start by preparing the knowledge graph data and historical cases for case based reasoning training. Subsequently, we also train all the five KGE model to generate embeddings for entities and relations.
- Prediction Generation: For the test set, for a given query (head entity, relation, tail entity), we run all the methods to generate prediction scores. For CBR methods, we retrieve similar historical cases and calculate a prediction score based on their outcomes or solutions. For KGE methods, we calculate the prediction score based on the distance between the embeddings of the head entity and the tail entity after applying the relation embedding. Hence, given a prediction models, we find prediction of a test query [drug→ treats→ disease]. Resulting in a list of prediction, which are a list of predicted disease each with a corresponding score. We consider score here, as list of reciprocal rank based on the position of each prediction in the list (Fig 2).

**Figure 2.**
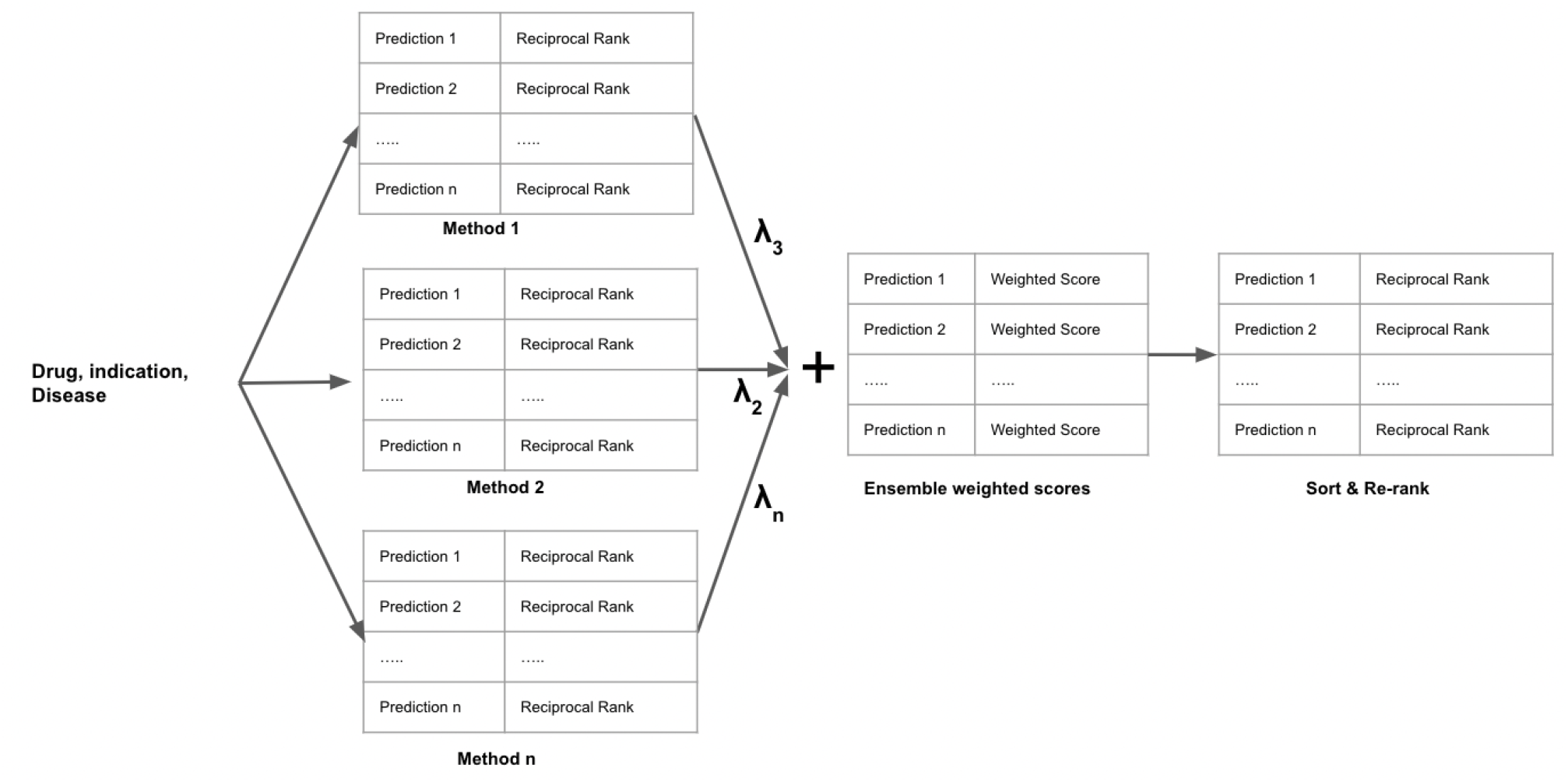
A more detailed representation of how we linearly combine the prediction score of various methods into a weighted ensemble method. Each prediction list is a list of all possible answers returned by a [Drug→ treats→ Disease] query and their corresponding rank of occurrence. The combined ensembled weighted score list is finally sorted with all the predictions re-ranked in this new list to give the final prediction of the WeightedKgBlend method
- Weighted Average Combination: We assign weights to the predictions generated by each methods as (*λ*_1_, *λ*_2_, *λ*_3_, *λ*_4_, *λ*_5_, *λ*_6_, *λ*_7_). We iterate each *λ* from range (0,1) and calculate a combined prediction score. The combination of *λ* which corresponds to the highest score is our final set of weights for all the methods. Formulating the scoring method, given a Knowledge Graph, G comprising of entities *v*_*i*_ ∈ V and relations *r*_*i*_ ∈ R, we define our hybrid scoring function P(e, G) as follows: *P*_*WeightedKgBlend*_(*e, G*) = *λ*_1_*· P*_1_(*e, G*) + *λ*_2_*· P*_2_(*e, G*) + *λ*_3_*· P*_3_(*e, G*) + *λ*_4_*· P*_4_(*e, G*) + *λ*_5_*· P*_5_(*e, G*) + *λ*_6_*· P*_6_(*e, G*) + *λ*_7_*· P*_7_(*e, G*), where: *P*_1_ : measure of an edge *e∈* [*v*_1_→ *r*→ *v*_2_] learned by TransE method *P*_2_ : measure of an edge *e∈* [*v*_1_→ *r*→ *v*_2_] learned by RotatE method *P*_3_ : measure of an edge *e∈* [*v*_1_→ *r*→ *v*_2_] learned by DistMult method *P*_4_ : measure of an edge *e∈* [*v*_1_→ *r*→ *v*_2_] learned by Rephetio method *P*_5_ : measure of an edge *e∈* [*v*_1_→ *r*→ *v*_2_] learned by ComplEx method *P*_6_ : measure of an edge *e∈* [*v*_1_→ *r*→ *v*_2_] learned by CBR method *P*_7_ : measure of an edge *e∈* [*v*_1_→ *r*→ *v*_2_] learned by Prob-CBR method *λ*_1_ : ensemble weight for TransE *λ*_2_ : ensemble weight for RotatE *λ*_3_ : ensemble weight for DistMult *λ*_4_ : ensemble weight for Rephetio *λ*_5_ : ensemble weight for ComplEx *λ*_6_ : ensemble weight for CBR *λ*_7_ : ensemble weight for Prob-CBR given: *λ*_1_ + *λ*_2_ + *λ*_3_ + *λ*_4_ + *λ*_5_ + *λ*_6_ + *λ*_7_ = 1
- Validation and Evaluation: Finally, we evaluate the final *λ* combination and the evaluate perfromance on the validation set to assess the ensembling method’s accuracy and effectiveness.
- Iterative Improvement: As we gather more data and improve the CBR system and KGE embeddings, we can continuously refine the weights and the combination strategy to achieve better results.

## Results

After iterating over all possible lambda combinations from (0, 1) condition that their summation is equal to one and evaluating the corresponding hybrid method on the test set, we found the lambda combination of (*λ*_1_ = 0, *λ*_2_ = 0.2, *λ*_3_ = 0, *λ*_4_ = 0.4, *λ*_5_ = 0, *λ*_6_ = 0, *λ*_7_ = 0.4) as the best performing MRR. Hence, we select this combination of lambda as our optimised weights for the WeightedKgBlend method. Table 1 shows the comparative benchmark of all the individual methods and WeightedKgBlend across various performance metric such as Hit@1, Hit@3, Hit@10 and mean reciprocal rank (MRR). Performance of WeightedKgBlend is higher across all metrics.

**Table 1.** Comparative Results.

| *Algorithm* | *MRR* | *Hits@1* | *Hits@3* | *Hits@10* |
| --- | --- | --- | --- | --- |
| *CBR* | 0.0424 | 0.00931 | 0.048 | 0.098 |
| *probCBR* | 0.1912 | 0.1157 | **0.3263** | 0.4526 |
| *TransE* | 0.1544 | 0.0 | 0.2421 | 0.4421 |
| *RotatE* | 0.1727 | 0.1052 | 0.1473 | 0.2526 |
| *DistMult* | 0.074168 | 0.017613 | 0.074364 | 0.176125 |
| *ComplEx* | 0.090426 | 0.027397 | 0.084149 | 0.232877 |
| *Rephetio* | 0.20228 | 0.1430 | 0.2657 | **0.4981** |
| **WeightedKgBlend** | **0.2387** | **0.1579** | 0.2442 | 0.37 |

We can see that from the nature of the equation, that WeightedKgBlend is assigning zero weight to the low performing algorithms like TransE, DistMult, ComplEx and simple CBR. Hence, we re-run the method but this time only considering the best three performing algorithms as observed by their assigned weights over the test dataset. We represent performance of the three best performing algorithm and the WeightedKgBlend method over all the lambda combination on the test set over several graph completion metric such as Hit@1, Hit@3, Hit@10 and MRR in Fig 3. Since, the four low performing methods are getting a zero assignment for our experiment, for simplicity we update the scoring method, given a Knowledge Graph, G comprising of entities *v*_*i*_ ∈ *V* and relations *r*_*i*_ ∈ *R*, we re-define the scoring function for WeightedKgBlend P(e, G) as follows:

*P*_*WeightedKgBlendTop*3_(*e, G*) = *λ*_1_*· P*_1_(*e, G*) + *λ*_2_*· P*_2_(*e, G*) + *λ*_3_*· P*_3_(*e, G*), where:

*P*_1_ : measure of an edge *e∈* [*v*_1_→ *r*→ *v*_2_] learned by RotatE method

*P*_2_ : measure of an edge *e∈* [*v*_1_→ *r*→ *v*_2_] learned by Rephetio method

*P*_3_ : measure of an edge *e∈* [*v*_1_→ *r*→ *v*_2_] learned by Prob-CBR method *λ*_1_ : embedding weight for RotatE

*λ*_2_ : embedding weight for Rephetio

*λ*_3_ : embedding weight for Prob-CBR

given: *λ*_1_ + *λ*_2_ + *λ*_3_ = 1

**Figure 3.**
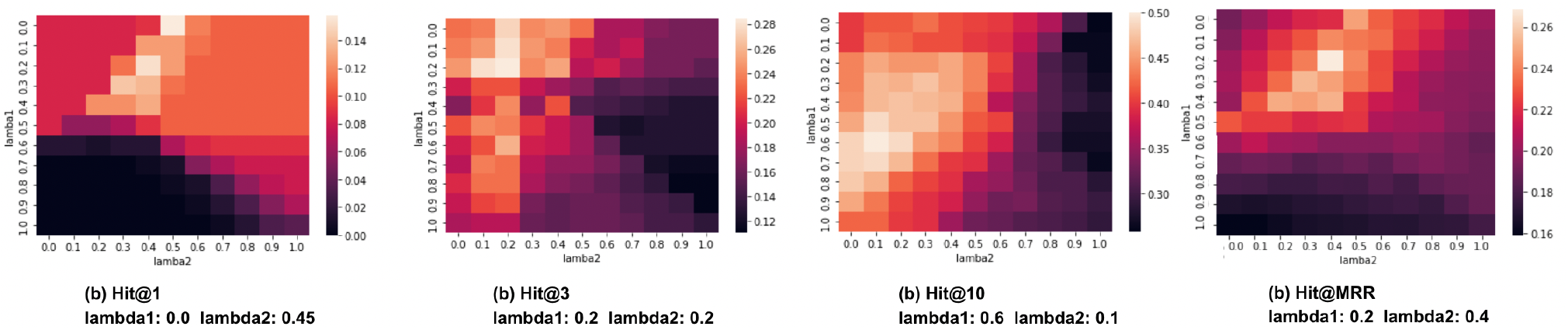
A heatmap for all the iteration of lambdas on the test set for top three methods (*λ*_1_: RotatE, *λ*_2_: Rephetio, *λ*_3_: Prob-CBR) over all the four metric: Hit@1, Hit@3, Hit@10 and MRR.

We represent the performance on the test set of WeightedKgBlend with the performance of the three best methods on the test set in Fig 4(a, b). Here, for WeightedKgBlend method we took the best lambda combination (*λ*_1_ = 0.2, *λ*_2_ = 0.4, *λ*_3_ = 0.4) as discussed before. We fix these lambda values as final optimized lambda and re-run the experiment on the valid test to measure an unbiased evaluation of the proposed method (Fig 5 (a, b)).

**Figure 4.**
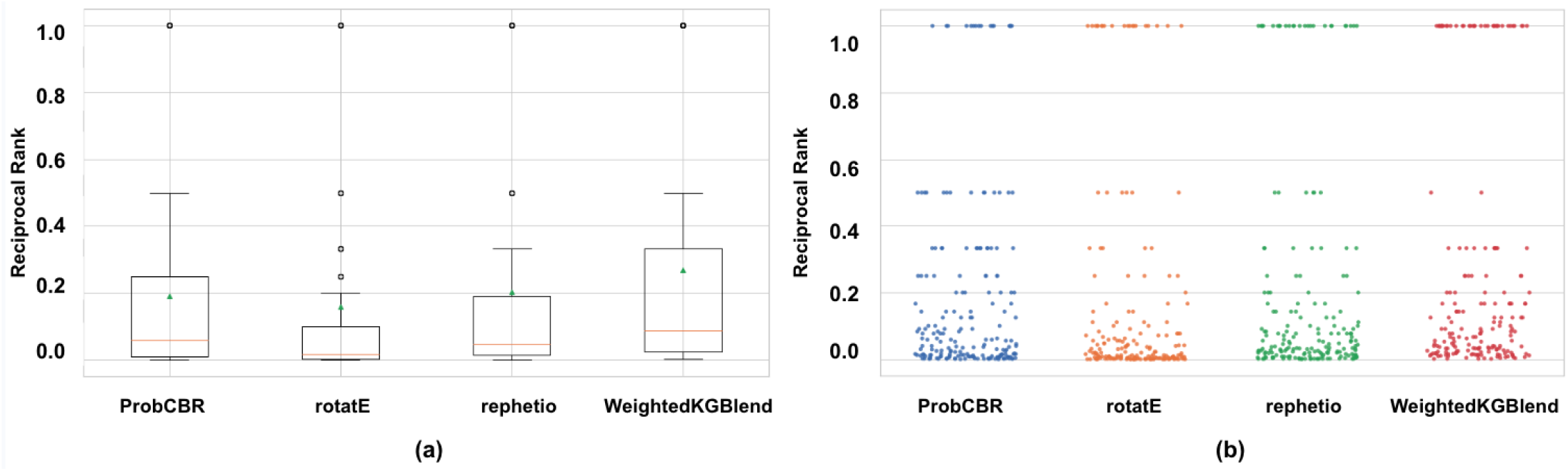
Box-plot (a) and Strip-plot (b) comparison of reciprocal ranks of the three best performing methods (RotatE, Rephetio and Prob-CBR) and WeightedKgBlend on the test set. The analysis suggests WeightedKgBlend outperforming the individual methods.

**Figure 5.**
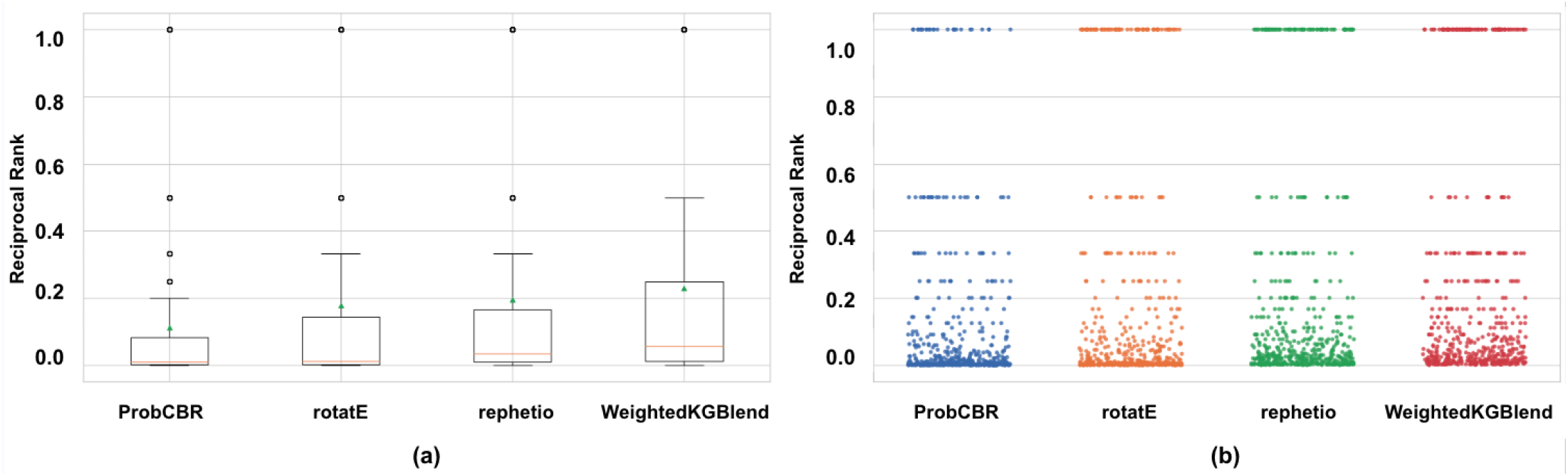
Box-plot (a) and Strip-plot (b) comparison of reciprocal ranks of the three best performing methods (RotatE, Rephetio and Prob-CBR) and WeightedKgBlend on the valid set. The analysis suggests WeightedKgBlend outperforming the individual methods.

Fig 6 represents an analysis of performance on some randomly selected [drug→ disease] examples from the test set. Here we show individual ranks of expected disease prediction by each models and WeightedKgBlend highlighted in green, where for better visualization more greener is higher rank. We notice for very few examples all the three individual methods agree on the ranking while for most examples there is a lack of consistency in rank. Probably due to algorithmic differences, some methods rank the expected answers higher while others do not. On ensembling these methods we find a better and more consistent performance in the WeightedKgBlend method. The KGE methods (RotatE and Rephetio) are more robust and scalable, while the case based reasoning method (Prob-CBR) is interpretable and gives us the path that lead to the correct answer as shown in the last column. Hence, combining these methods not only gives us a better performance but also makes the prediction more interpretable.

**Figure 6.**
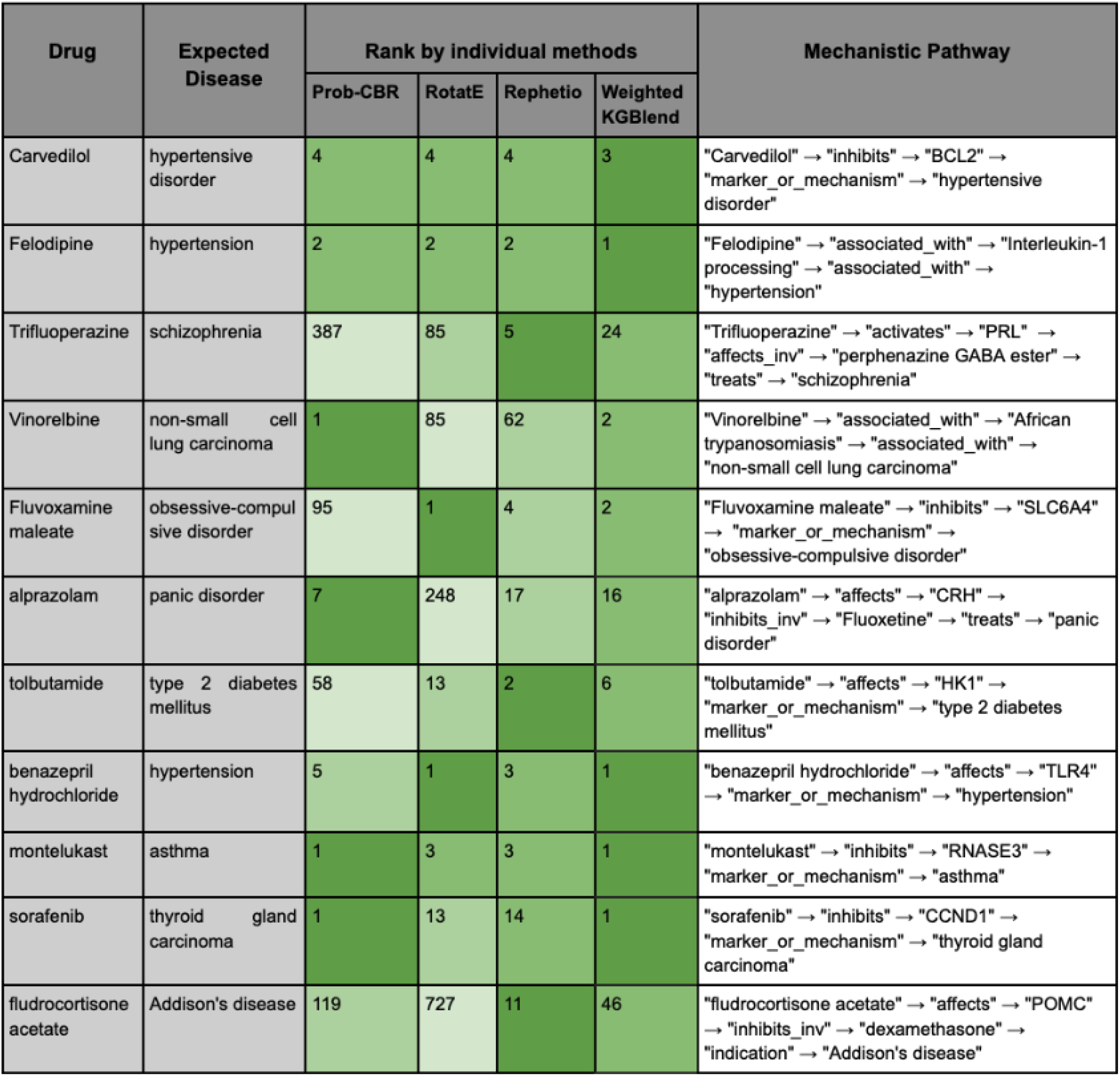
Few examples of prediction along with their respective rank of the expected answer by each comparative method. More greener is a higher rank.

### 0.4 Drug repurposable cases

We selected a few examples where we verify the correctness of the topmost predicted disease for a drug by our method to study the potential identification of drug repurposable candidates.

- Carvedilol → indication → Acute coronary syndrome The topmost predicted disease by WeightedKgBlend by drug “Carvedilol” is “Acute coronary syndrome”. Carvedilol is primarily used in the management of chronic heart failure and hypertension, as well as to reduce the risk of death after a heart attack (myocardial infarction). While carvedilol is not commonly used as a first-line treatment for acute coronary syndrome (ACS), there is some ongoing research investigating its potential repurposing or adjunctive use in ACS management^20^. Carvedilol belongs to a class of medications known as beta-blockers, which work by blocking the effects of certain stress hormones such as adrenaline on the heart and blood vessels^21^. Some studies have suggested that carvedilol may have benefits in ACS beyond its primary indications. For example, it may help reduce myocardial ischemia (lack of blood flow to the heart muscle) by decreasing heart rate and blood pressure, thereby reducing the workload on the heart. Additionally, it has antioxidant properties that could potentially offer cardioprotective effects during ACS.
- Fluvoxamine maleate → indication → major depressive disorder The topmost predicted disease by WeightedKgBlend by drug “Fluvoxamine maleate” is “major depressive disorder”. Fluvoxamine is an approved medication for the treatment of obsessive compulsive disorder in adults and children ages 8 and older. Fluvoxamine may also be helpful when prescribed “off-label” for major depressive disorder, social phobia (also known as social anxiety disorder), posttraumatic stress disorder, panic disorder, and eating disorders including bulimia nervosa and binge-eating disorder^22^. Fluvoxamine is approved for treating depression in the UK but not in the U.S.
- Tolbutamide → indication → breast cancer The topmost predicted disease by WeightedKgBlend by drug “Tolbutamide” is “breast cancer”. Tolbutamide is used to treat high blood sugar levels caused by a type of diabetes mellitus (sugar diabetes) called type 2 diabetes. Several retrospective observational studies found that patients treated with tolbutamide had lower cancer mortality than patients treated with glibenclamide. Some studies have shown that antidiabetic drugs can inhibit the progression of cancer cells, but others have shown that they can also be a risk factor in cancer development or progression. A 2024 meta-analysis found no significant association between the use of TZDs and the risk of breast cancer in randomized controlled trials and case–control studies. However, a more recent meta-analysis found that the use of thiazolidinediones was associated with a lower risk of breast cancer.

## Conclusion

This study uses the synergy potential of knowledge graph embedded (KGE) and case-based reasoning (CBR) methods to develop a new drug discovery methodology using a weighted ensemble method. In this work, we proposed a framework that dynamically integrates predictions of various KGE and CBR models and assigns weights according to individual performance. Our results show that the Weighted Ensemble Method exceeds independent approaches, achieving greater predictive accuracy and robustness. Our integration method utilizes KGE’s complementary strengths in capturing complex relationships unstructured data and in drawing parallels from historical examples to provide a comprehensive and nuanced understanding of complex biomedical landscapes.This research highlight the transformational potential of the integration of various methods of artificial intelligence to enhance the discovery of biomedical knowledge as well as shows advances the field of drug reuse.

## Author contributions statement

MS wrote the main manuscript. MS implemented the method with help of previous and current work of RT and CG. AIS helped with mentoring and reviewing the work.

